# Ancient microRNA profiles of a 14,300-year-old canid are taxonomically informative and give glimpses into gene regulation from the Pleistocene

**DOI:** 10.1101/2019.12.16.877761

**Authors:** Bastian Fromm, Marcel Tarbier, Oliver Smith, Love Dalén, M. T. P. Gilbert, Marc R. Friedländer

## Abstract

Ancient DNA sequencing is the key technology for paleogenomic studies and today a routine method in many laboratories. Recent analyses have shown that, under favoring conditions, also RNA can be sequenced from historical and even ancient samples. We have re-analyzed ancient RNA data from a Pleistocene canid and find - in addition to the previously described messenger RNA fragments - intact microRNAs, which are short transcripts with important gene regulatory functions. With an extraordinary age of 14,300 years, the canid microRNA profiles are the oldest ever reported. Despite their age, we show that the microRNA profiles are conclusive of taxonomic origin, tissue identity with organ- and cell-type specific signatures, and that they yield glimpses into gene regulatory activity and biological processes from the Pleistocene. In summary, we here show that straightforward microRNA analyses hold great promise for deeper insights into gene regulation in extinct animals.

## Introduction

Sequencing historic and ancient DNA from up to 600,000-year-old samples has become a standard approach to infer the genetic history of extinct or extant species (Orlando et al. 2013). While much has been learned from studying these genomes, it remains difficult to infer cellular processes, such as in vivo genome function, from DNA directly.

In contrast, RNA conveys genetic information at the functional level, and can thus give insights into the biological activity, the tissue identity, or even the cellular composition of samples (Newman et al. 2015). However, based on a putative relative lack of surviving material in historic and ancient samples, due to the release of RNases during decomposition in most tissues (Huynen et al. 2012), ‘ancient RNA’ (aRNA) has rarely been studied and is often disregarded as a source for biological discoveries. On the other hand, preservative conditions for aDNA (cold, draught) actually also inhibit RNAses, and there are significantly more RNA molecules than DNA found per cell (see for microRNA (Calabrese et al. 2007)).

With the availability of powerful and sensitive sequencing approaches, a small number of recent studies have capitalised on early observations of very short RNA fragments surviving in some archaeological plant and mummified materials (Rollo 1985; Venanzi and Rollo 1990), and have recovered sequenceable amounts of aRNA from historic plant and feces samples. In doing so they have detected, for instance, viral RNA genomes (Ng et al. 2014; Smith et al. 2014), microRNAs (Smith et al. 2017) and even fragments of protein-coding transcripts (Fordyce et al. 2013). While these studies represent a proof of concept for aRNA sequencing, or paleotranscriptomics, they are limited to relatively recent (<1000 years old) samples and, except for one study, restricted on seed-material that conveys highly preservative conditions.

A recent sequencing study on three permafrost and two historical canid samples showed that aRNA can survive for extended periods in mammalian samples (Smith et al. 2019). With approximately 14,300 years of age for the Pleistocene permafrost samples, the authors presented the oldest ever sequenced RNA to date. In a paleotranscriptomic analysis, using two custom bioinformatics methods based on total aRNA profile comparisons to recent tissue sample profiles, aRNA profiles could resolve tissue identity for two of the samples.

Because the applied sequencing strategy (single-end, short read, without size-selection) clearly favors short RNAs, we here investigate whether microRNAs can also be identified in the data. MicroRNAs are 22 nucleotide short RNA molecules that are excellent cell- and tissue markers (Christodoulou et al. 2010; McCall et al. 2017; de Rie et al. 2017), and stable in historic samples (Keller et al. 2017; Smith et al. 2017). They are produced when RNA hairpin structures (‘precursor microRNAs’) are cleaved into functional regulatory molecules (‘mature microRNAs’) and biogenesis by-products (‘star microRNAs’). The mature microRNAs are key regulators of protein coding genes with important functions in numerous biological processes including development and disease (Bartel 2018). They are the most conserved elements in metazoan genomes and have great potential as phylogenetic markers (Sempere et al. 2006; Tarver et al. 2013). We have recently curated the microRNA complements of 45 Metazoan organisms, including canid, in the MirGeneDB database (Fromm et al. 2019) and shown that species-specific microRNAs can be used to trace the taxonomic origin of samples using our straightforward software miRTrace (Kang et al. 2018). We here leverage these two resources to characterize the microRNA complements of the ancient and historic canid samples.

## Results

### Abundant microRNA detection in ancient and historic samples

Reanalyzing the RNAseq datasets (Illumina) for the ancient and historic samples from Smith et al. (Smith et al. 2019), we detected 334 microRNAs (236 mature and 98 star) out of the 447 currently annotated canid microRNAs (Fromm et al. 2019) (Figure 1A), with in total more than 18,000 sequence read-outs, or ‘reads’, among all samples (Figure 1B). Interestingly, the ratio between detected distinct ‘mature’ and ‘star’ products was relatively constant between samples (ranging from 2.3 in ancient liver to 3.0 historic skin samples), suggesting that the two types of molecules are equally affected by time, when preserved in permafrost. Analyses on previously reported damage patterns on aRNA showed comparable patterns in microRNAs (Jónsson et al. 2013; Smith et al. 2019) (Supplementary Figure 1). The overall high number of detected mature and star microRNAs and reads, representing 50% of all currently known canid microRNAs, was highest for the historic samples and lower for the ancient samples (Figure 1B and 1C). Among the ancient samples, the liver showed the best preservation, which is in line with experimentally observed postmortem migration patterns of bacteria, that showed near-sterile conditions in liver up to 5 days postmortem, which clearly is a reasonable timescale for the Tumat puppy to be frozen (Tuomisto et al. 2013).

**Figure 1:**
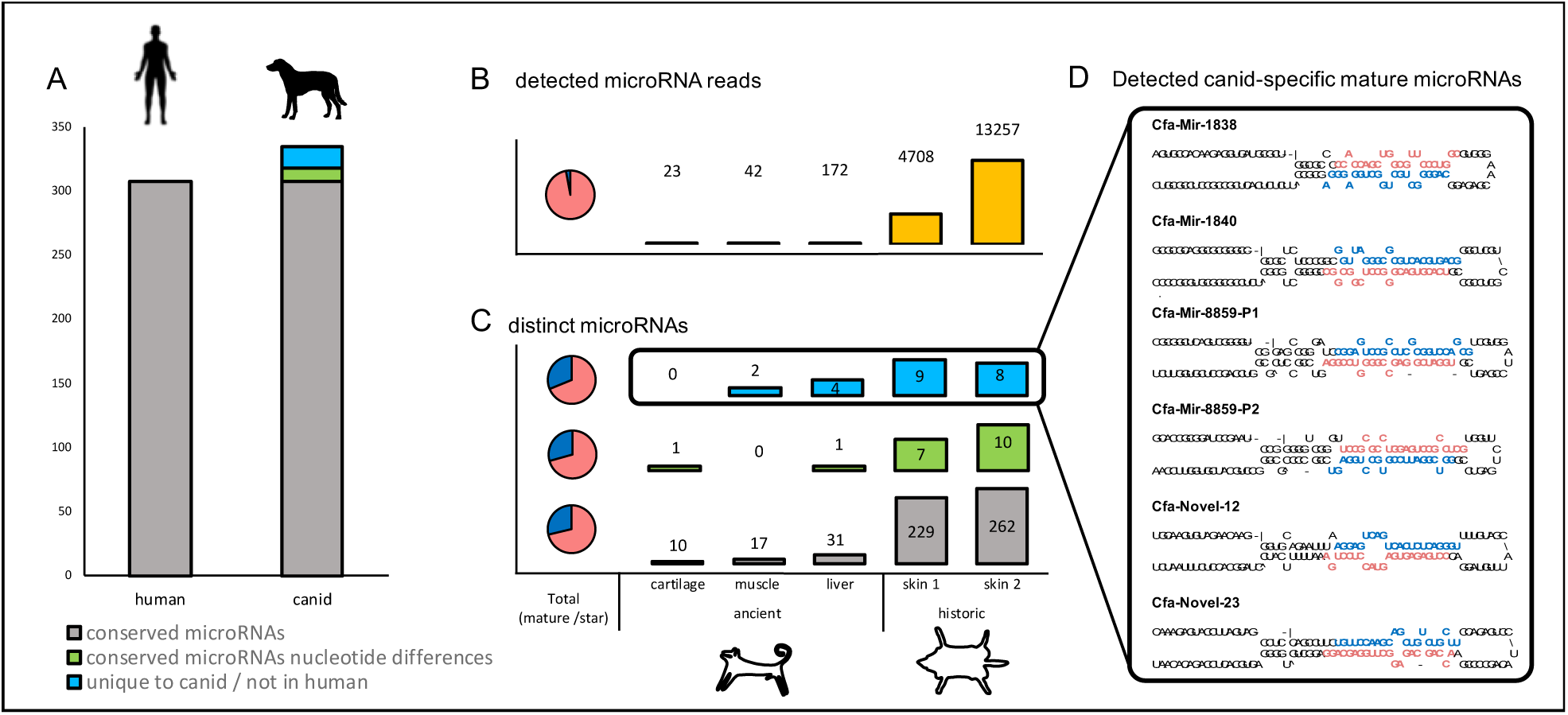
Abundant numbers of conserved and dog-specific microRNAs are detected in historic and ancient samples. A) 334 microRNAs (236 mature and 98 star) are detected in the historic and dog samples, of those 307 microRNAs (219 mature and 88 star) are identical in sequence to human (grey), 11 microRNAs (6 mature and 5 star) show single nucleotide differences (polymorphic) (green) and 16 microRNAs (11 mature and 5 star) are absent in human (blue). B) Total number of microRNA reads detected in each sample (yellow). C) Detailed numbers of conserved, polymorphic and canid-specific microRNAs detected in the 5 ancient and historic samples. D) Hairpin structures and indication of mature (red) and star (blue) products of 6 of the 11 microRNAs not found in human that are specific to canid and currently not known in any other organism.

### microRNAs support taxonomic origin of the sequences

Given the extreme age of the canid aRNA samples, it is a concern that the detected sequences do not originate from the ancient tissues, but rather from trace levels of contamination. Using a large range of species- and clade-specific microRNAs as reference, our software miRTrace did not identify any microRNA-based contamination by non-canid eukaryotes (Supplementary File 1). However, during evolution, the majority of microRNAs are highly conserved in sequence (Fromm et al. 2015; Bartel 2018), and it is important to investigate if the sequences of the detected microRNAs can be used to unambiguously identify the taxonomic species of a sample. The accurate annotations of MirGeneDB enabled us to make sequence comparisons for each microRNA to human and canid, and test for conservation and variation in the 334 individual microRNAs (236 mature and 98 star).

As expected, we found that the majority of detected microRNAs, in total 307 microRNAs (219 mature and 88 star) belongs to microRNAs that are conserved between human and canid. Thus, these microRNAs are not informative for species delimitation, and, at least theoretically, could derive from contaminations. This is, however, unlikely given the absence of any microRNAs specific to other clades, as shown in the miRTrace analysis.

A canid origin of the sequences was supported by 11 distinct microRNAs (6 mature and 5 star) that showed clear nucleotide differences to their orthologues in human (‘polymorphic microRNAs’). The six mature microRNAs also showed differences to other organisms, none of which can be explained by deamination events (C->U conversions; Mir-8-P3a, Mir-28-P2, Mir-154-P2, Mir-339, Mir-193-P1b and Mir-105-P2; Supplementary File 2, Supplementary Figure 2B).

Finally, we found 16 microRNAs (11 mature and 5 star) that are present in canid but completely absent in human (‘microRNAs unique to canid’). Importantly, 6 of the 11 mature and 3 of the 5 star microRNAs were also not found in other MirGeneDB species and appears to be truly canid-specific, having never evolved in any other animal group (Mir-8859-P1 and P2, Mir-Novel-23, Mir-Novel-12, Mir-1838 and Mir-1840; Figure 1D). This not only renders them candidate biomarkers for the authenticity of the sample (Figure 1C, D), but also makes them stand out as interesting regulatory molecules with putative canid-specific functions. The remaining seven microRNAs (5 mature, 2 stars) were not found in human, but were found in other organisms. Of those, single nucleotide differences to at least one or several other species orthologous sequences were found in five microRNAs (Mir-1842, Mir-154-P34, Mir-1388, Mir-2387, and Mir-7180; Supplementary Figure 2A), making them informative for a distinction with canid, too.

When comparing the levels of detected conserved, polymorphic and unique microRNAs, we saw that not only the historic samples, but also the ancient samples showed all three categories of microRNAs, supporting the authenticity of the microRNA reads (Figure 1C). Altogether, the identification of canid-specific microRNAs and canid-specific microRNA variants strongly supports the authenticity of the sequencing data and provides evidence that no contaminations from closely related species confound our results.

### microRNAs inform about cellular and tissue identity of ancient and historic samples

For extant organisms, it was shown that a number of microRNAs have clear tissue- and sometimes even cell-type specific expression patterns (Christodoulou et al. 2010; McCall et al. 2017; de Rie et al. 2017). In addition, using involved and custom methods, Smith et al. 2019 demonstrated that total aRNA could be used to confirm the identity of ancient liver and one of the historic skin samples (Smith et al. 2019). Therefore, we next asked whether the tissue identity of the samples could be inferred based on the microRNAs as well.

When looking at the top 5 expressed microRNAs of each sample alone, clear signals of tissue and cell-type specific microRNAs were identified. Specifically, we identified Mir-205-P1, a skin and cartilage specific microRNA, as the microRNA with highest abundance in both skin samples and ancient cartilage and Mir-203, with similar tissue specificity, in both historic skin samples (Teta et al. 2012). The ‘myoMir’ Mir-133 (Sempere et al. 2004) was found among the top 5 microRNAs in the ancient muscle sample, and the hepatocyte-specific Mir-122 (McCall et al. 2017) was among the top 5 microRNAs in the ancient liver, again confirming the identity of these tissues using microRNAs as markers.

Interestingly, we also noticed relatively high levels of microRNAs that are not strictly specific to any of the tissues (cartilage, muscle, liver and skin), but are known to be specific to range of immune-cells (Table 1). Most notably, the lymphocyte-specific microRNA Mir-155 was detected in all samples and the highest overall detected microRNA in ancient muscle. Mir-148-P1, which is mast-cell specific was detected in all but ancient muscle samples and highest in the ancient liver sample. The high level of these microRNAs suggests a high number of immune-cells in the samples.

**Table 1:** The relative rank of immune-cell specific microRNAs detected in ancient and historic canid samples. Average ranks were computed for each sample independently. The lower the number, the higher the rank.

| microRNA | cell-type | aCa | aMu | aLi | hS1 | hS2 |
| --- | --- | --- | --- | --- | --- | --- |
| Mir-155 | Lymphocytes | 7 | 1 | 37 | 77 | 67 |
| Mir-339 | Macrophages | - | - | 7 | 45 | 19 |
| Mir-148-P1 | Mast-cells | 7 | - | 1 | 6 | 14 |
| Mir-378 | Dendritic cells | - | 3 | 2 | 3 | 6 |
| Mir-24 | Macrophages | - | 3 | - | 8 | 6 |

Next, we compared the five samples with more than 69 smallRNAseq datasets of fresh soft dog tissues from two previously published studies (Koenig et al. 2016; Penso-Dolfin et al. 2016) using MirGeneDB 2.0 annotations (Fromm et al. 2019). In a principal component analysis (PCA), based on the normalized counts of microRNA sequence reads only, we found that recent samples cluster well according to tissue and organ group (Figure 2). Two of the ancient samples, ‘ancient cartilage’ and ‘ancient muscle’, which were previously also shown to have the weakest tissue identity signal based on a customized total aRNA analyses (Smith et al. 2019), showed less than 20 expressed distinct microRNAs, with less than 100 microRNA reads in total and did not yield enough information to be included in the PCA. However, the ancient liver sample and the two historic skin samples, clustered well with recent liver and skin samples, respectively (Figure 2).

**Figure 2:**
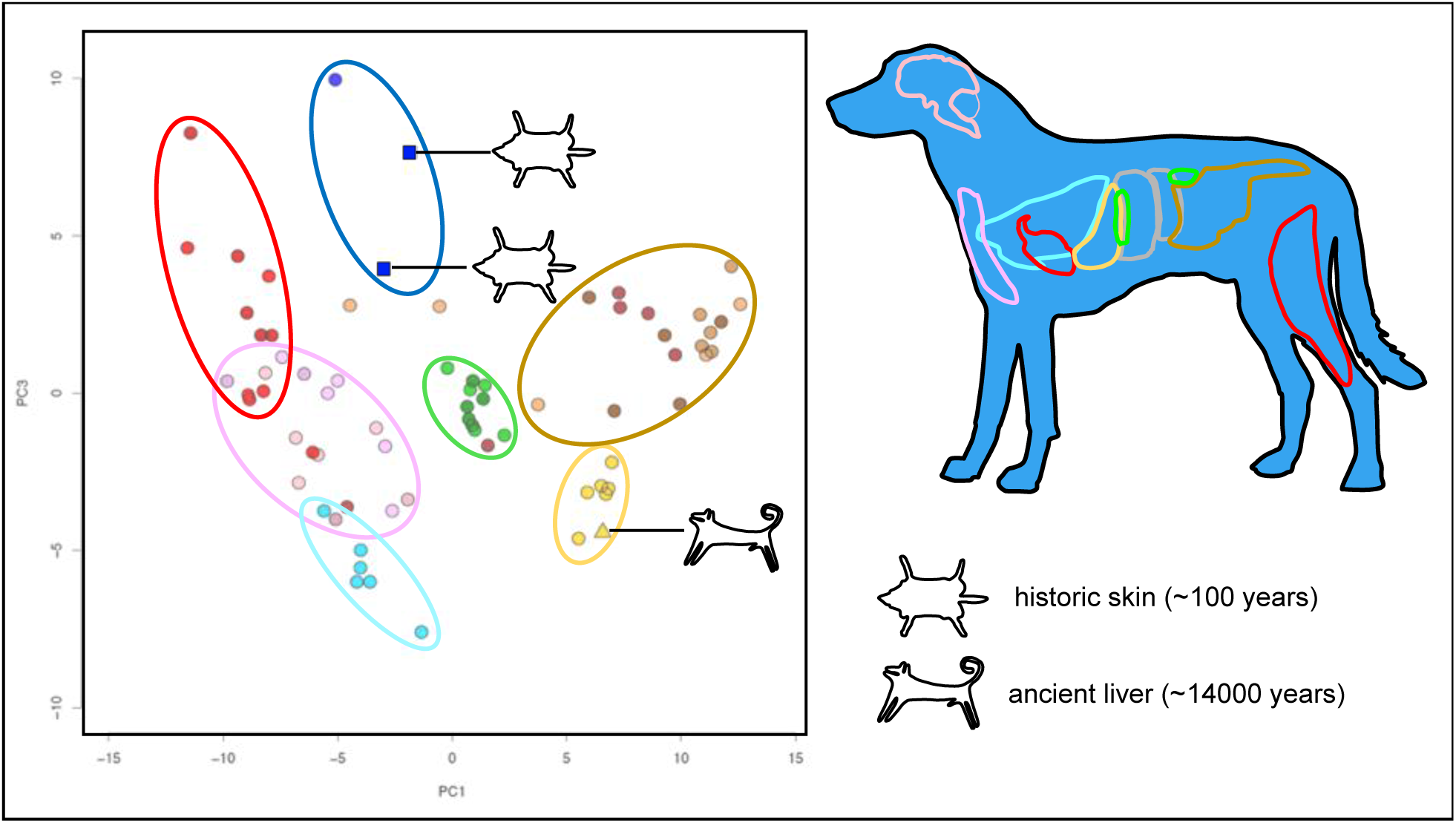
MicroRNA profiles of ancient and historic samples resemble recent tissue expression and are informative for tissue identity. PCA plot of in total 72 samples (69 recent, 1 ancient, 2 historic) show clustering of major organ groups (red - muscle/heart, pink - thymus and brain, light blue - lung, green - kidney and pancreas, dark blue - skin, brown - gastrointestinal tract, yellow - liver). Historic skin samples (dark blue squares) cluster with recent skin samples and the ancient liver sample (yellow triangle) clusters with recent liver samples.

### A glimpse into ancient genome activity through microRNAs

As microRNA themselves are active regulatory molecules, their presence and differential abundance in different tissues alone can directly provide insights into gene regulatory activity (Bartel 2018). Indeed, when we studied the annotated functions of the predicted canid mRNA targets of the tissue-specific microRNAs, we found enrichment for functional annotations corresponding to biological processes clearly related to the respective tissue origin. For instance, the predicted targets of Mir-203, which is highly abundant in the historic skin samples, are enriched for the gene ontology “Epithelium and blood vessel development”, which is clearly skin related. Similarly, were the targets of Mir-122, which is abundant in ancient liver samples, enriched for “carbohydrate metabolism” and “response to starvation”, representing well-established liver functions. Ancient cartilage-specific Mir-205-P1 targets genes that are enriched for “Platelet-derived growth factor receptor”, which has been described to regulate chondrocyte proliferation (Kieswetter et al. 1997), and the ancient muscle specific Mir-133 targets genes enriched for ontologies related to muscle contraction (“Divalent inorganic cation transmembrane transporter activity”) and neuro-transmitters (“GABA receptor binding”). A full list of all specific microRNA and the enrichment of functional annotations within their target gene populations can be found in Supplementary Table 1. The intriguing distribution of some cell-type specific microRNAs across all samples can be found in Supplementary Figure 3.

## Discussion

With an estimated age of 14,300 years, we here present the oldest microRNA sequences to date. We report an abundant variety of microRNAs in three ancient and two historic samples and find that the microRNA profiles are informative and conclusive of taxonomic origin for all samples. We further show their ability to identify not only tissue-origin of all samples, but also, to some extent, the likely cellular composition of our samples, including an intriguing signal for high numbers of immune-cells in the sample. These findings encouraged us to ask more functional questions that might help us to get insights or glimpses into the *in vivo* genome activity of extinct animals. By predicting the targets of the top microRNAs in the samples, we could indeed identify pathways and cellular processes that the samples were likely conducting, and the microRNAs were regulating, 14,300 years ago.

In comparison to analyses of ancient mRNA, microRNA analysis is straight-forward, requiring only simple sequence matching that can be performed without mapping to a reference genome. Further, because microRNAs are captured with protocols that are optimized for highly degraded and fragmented RNA, microRNA analysis could be used as an alternative paleotranscriptomic approach to aRNA data in general. Importantly, since both mature and star sequences are detected well, it would in principle be possible to reliably predict microRNAs *de novo* in extinct species, possibly discovering novel microRNA families that are not present in any living animals (Ambros et al. 2003; Friedländer et al. 2008; Fromm 2016).

The detection of abundant numbers of intact microRNAs in pleistocene permafrost samples represents a proof of concept and opens up novel opportunities for future studies on the *in vivo* genome activity in other ancient samples, in particular those from extinct animals such as mammoths, cave lions and others that the increasingly melting permafrost might provide.

## Materials and Methods

Briefly, Illumina samples from ancient and historic (Smith et al. 2019) as well as fresh tissues (Koenig et al. 2016; Penso-Dolfin et al. 2016) were downloaded from SRA using sra-toolkit 2.9.2 (Leinonen et al. 2010), processed using miRTrace (Kang et al. 2018) and quantified using the ‘quantify’ function of miRDeep2 (Friedländer et al. 2012) with the MirGeneDB 2.0 dog microRNA complement as reference (Fromm et al. 2019). Sequence comparisons to human and other vertebrate microRNA complements are based on MirGeneDB 2.0 annotations (Fromm et al. 2019) and the consistent nomenclature of microRNA gene orthologues, paralogues and families (Fromm et al. 2015). Alignments were checked using custom scripts and AliView alignment viewer (Larsson 2014).

PCA analyses were based on ubiquitously expressed microRNAs (more than 95% of all samples, and ancient liver) on all tissue samples, excluding ancient cartilage and ancient muscle, bone marrow and sexual organs such as testis and ovary.

Tissue-specific microRNA were defined as those that are abundant in the tissues of interest but not abundant in other tissues included in this study. Targets of these microRNA were downloaded from TargetScanHuman using the lift over to dog transcripts. Gene ontology and KEGG pathway enrichment analysis were performed on the top 400 targets of these microRNA using the R packages topGO and clusterProfiler. Relevant annotations were selected from the (up to) 25 highest ranking significant terms.

## Supplementary Figures

**Supplementary Figure 1:**
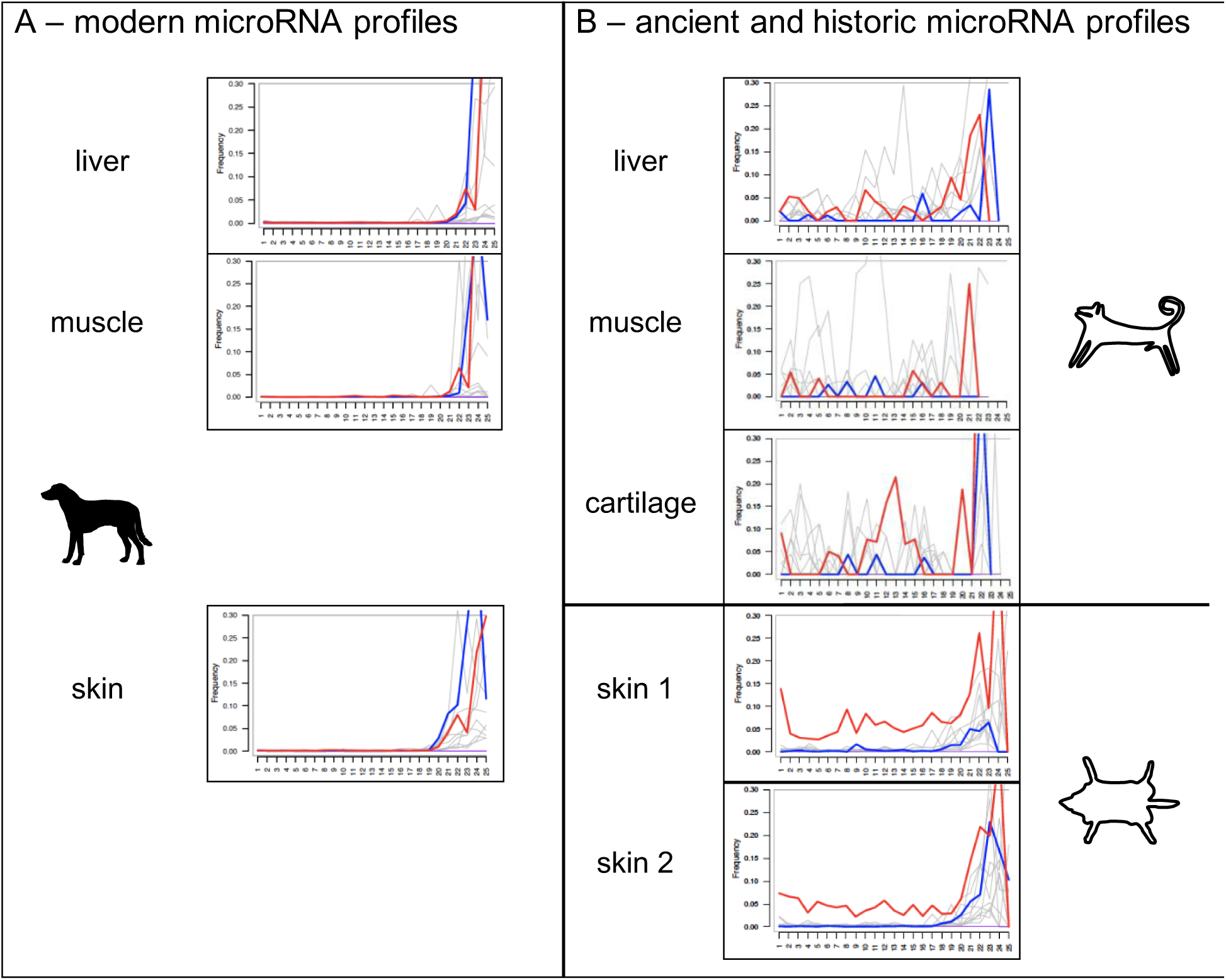
mapDamage profiles of A) modern and B) ancient and historic canid tissue samples to canid microRNA loci. Red lines show C>U transitions, while blue lines show G>A transitions. Note that the observation of 3’ non-templated nucleotides, in particular uridylations, is a hallmark of microRNAs. For the historic samples, overall conversion rate appears to be comparable to those described for ancient mRNA. However, for microRNAs, deaminase events appear to be the most frequent conversion type by far. For the ancient samples, nucleotide conversion rates are high and do not appear to be highly enriched for deaminase events. However, given the relatively low number of microRNA reads detected, we are reluctant to make strong conclusions at this point.

**Supplementary Figure 2:**
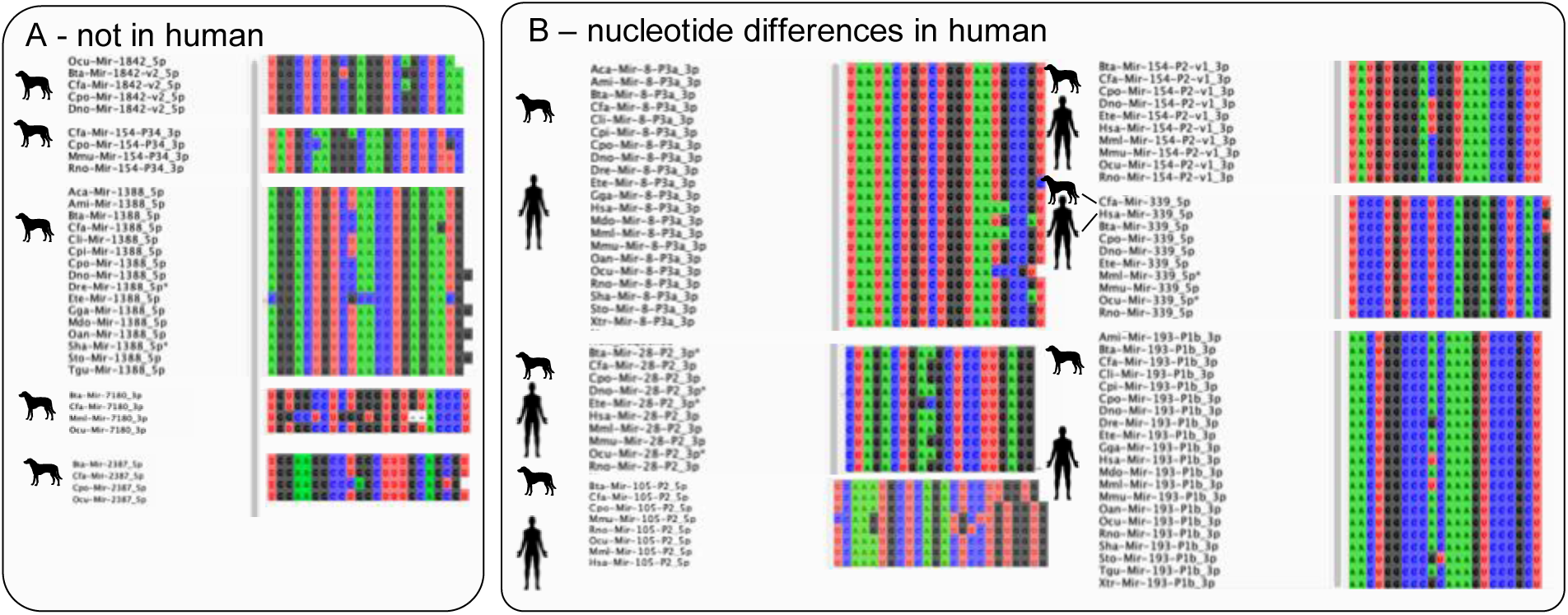
Conserved microRNAs that are taxonomically informative because they are either A) not found in human or B) found in human and showing nucleotide differences to at least one other MirGeneDB organism or human, respectively.

**Supplementary Figure 3:**
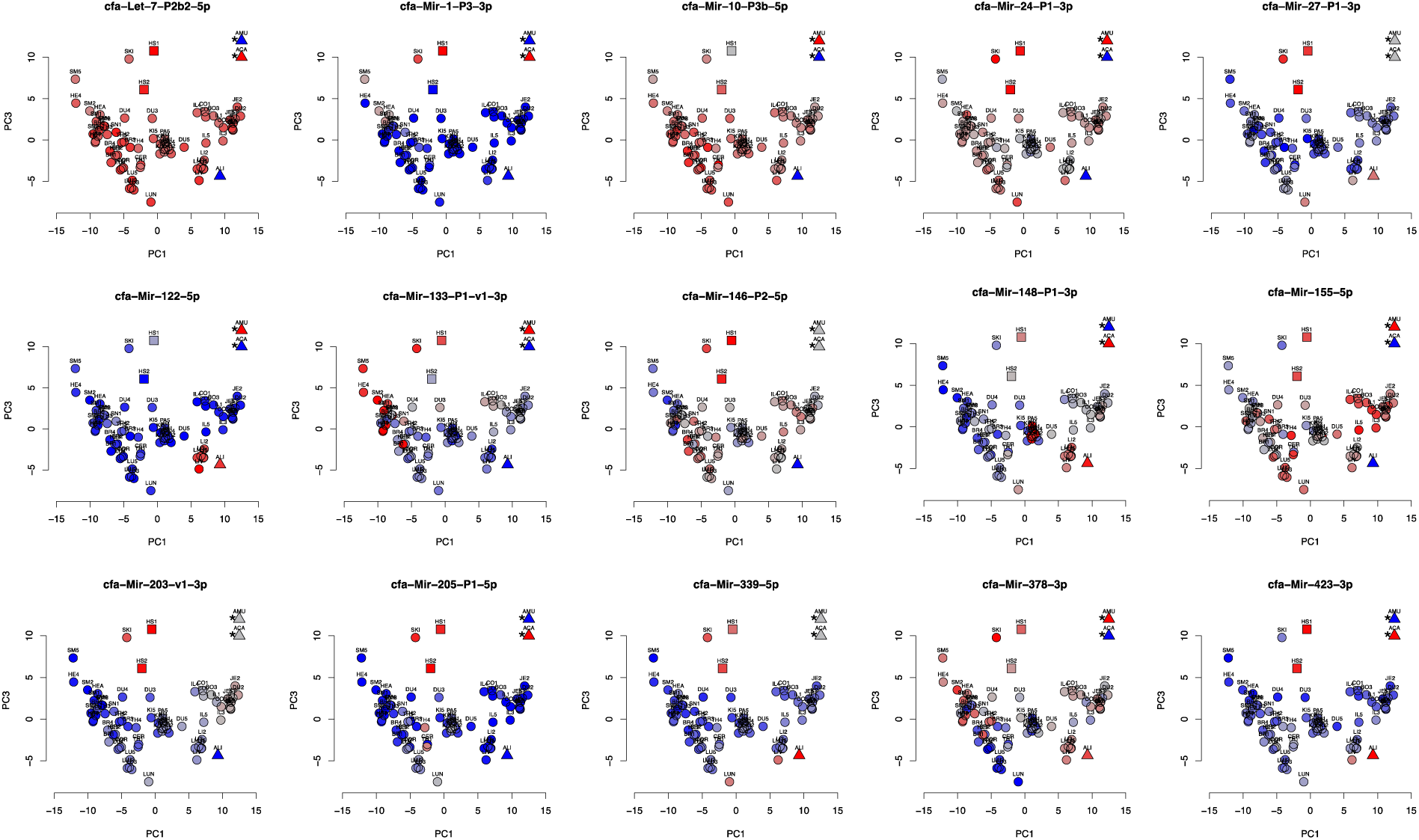
Cell type- and Tissue-Specificity of selected microRNAs across 69, 3 ancient and 2 historic samples. The PCA plot from Figure 2 was re-used to depict the color-coded detection of 15 selected microRNAs (blue - low, red -high), note that ancient cartilage and muscle samples were included to show expression of microRNAs. Thus, their relative position does not depict their distance to the other samples.

Supplementary Table 1 – Summarized gene ontology enrichments of targets of 1a ancient liver specific microRNAs, 1b ancient muscle specific microRNAs, 1c ancient cartilage specific microRNA, and 1d historic skin specific microRNA

**Supplementary Table 1a.** Summarized gene ontology enrichments of targets of ancient liver specific microRNA

| microRNA | Enriched terms |
| --- | --- |
| cfa-Mir-122-5p | Carbohydrate metabolism (GO:0005996, 0005975, 0019318, ...)<br>Response to Starvation (GO:0042594, 0042149, 0009267)<br>Growth factors (GO:1990090, 1990089, 0070848, ...) |
|  | Transcriptional regulation (GO:0000978, 0001067, 0000977, ...) |
| cfa-Mir-339-5p | Estrogen and androgen metabolism (GO:0006703, 0008209, 0008210, ...)<br>Fatty acid and CoA biosynthesis (GO:0046949, 0035337, 0035384, ...) |
|  | Enzymatic activity (GO:0047025, 0047035, 0003857, ...) |
| cfa-Mir-423-3p | Protein deglycosylation (GO:1904380, 0036507, 0036508, ...)<br>Misfolded protein reaction (GO:0071712, 0071218, 0051788, ...) |
|  | Transcriptional regulation (GO:0043565, 0003682, 1990837, ...) |
|  | Hippo signaling pathway (hsa04390) |

**Supplementary Table 1b.**
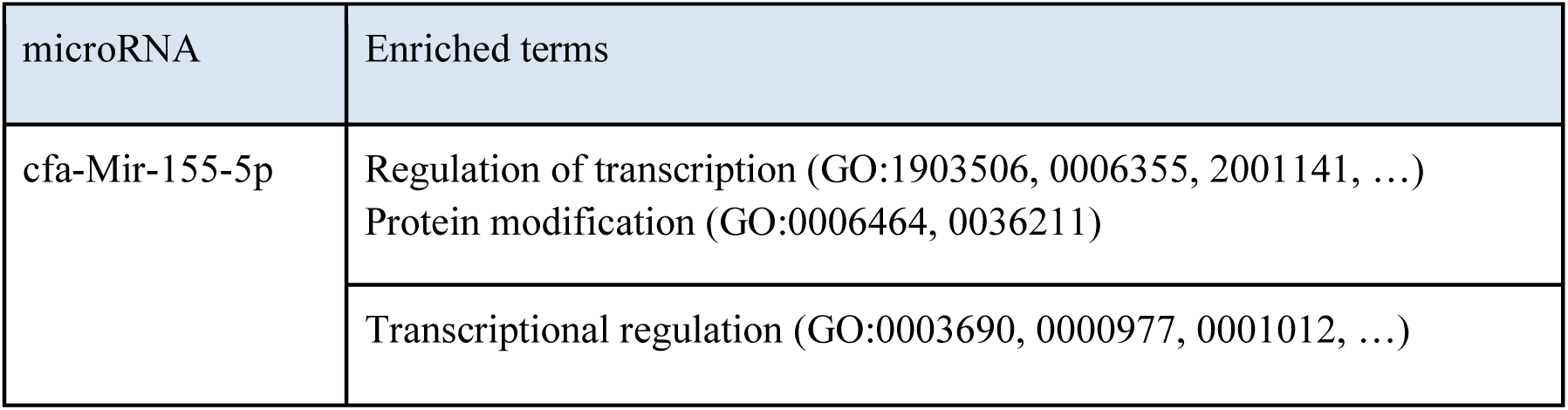
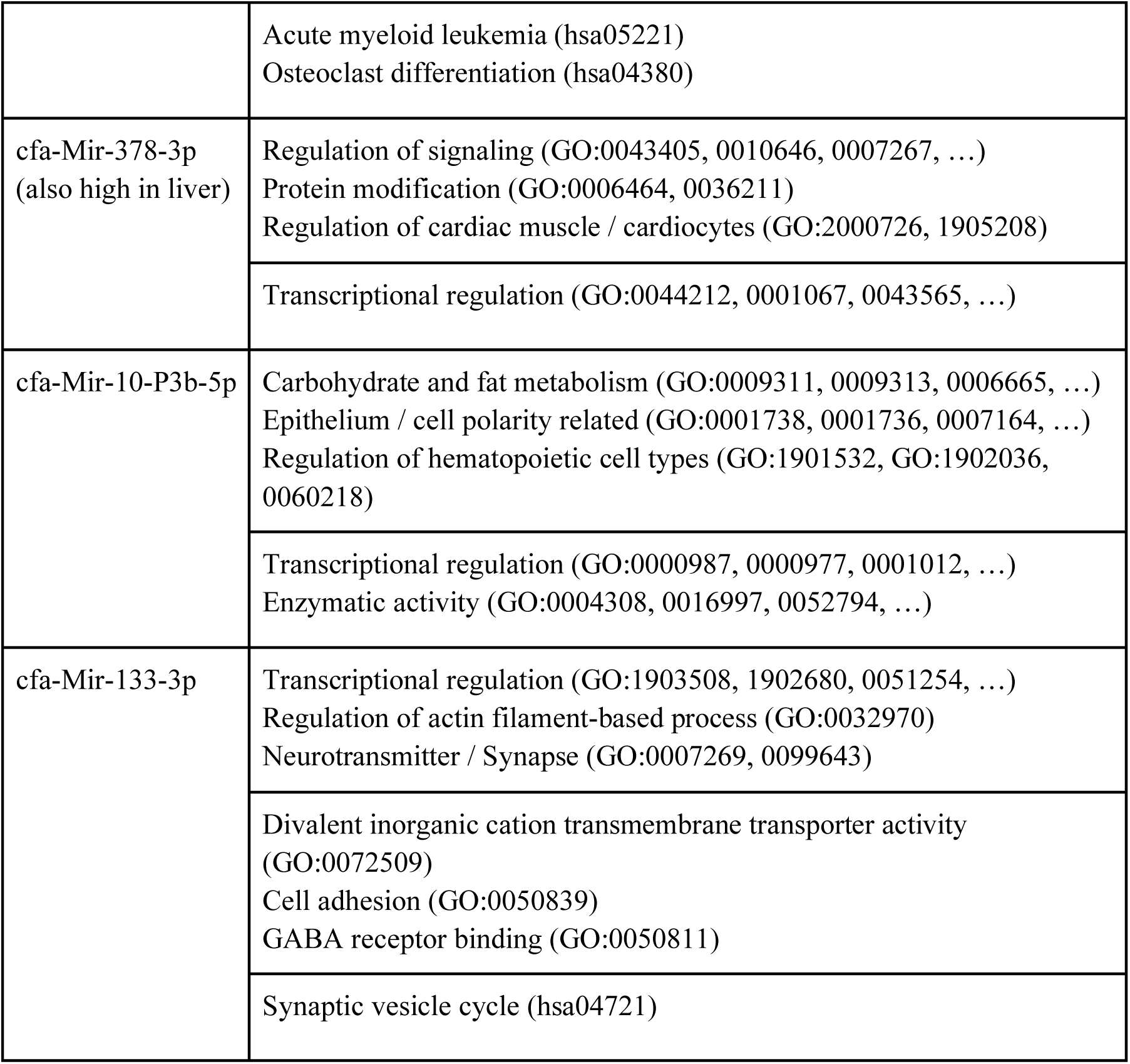
Summarized gene ontology enrichments of targets of ancient muscle specific microRNA

**Supplementary Table 1c.** Summarized gene ontology enrichments of targets of ancient cartilage specific microRNA

| microRNA | Enriched terms |
| --- | --- |
| cfa-Mir-205-P1-5p | Regulation of signaling (GO:0023051, 0010646, 0023056, ...) |
|  | Transcriptional regulation (GO:1990837, 0003690, 0043565, ...)<br>Enzyme activity (GO:0004308, 0016997, 0052794, ...)<br>Platelet-derived growth factor receptor (GO:0005161) |
|  | Hippo signaling pathway (hsa04390, hsa04392) |
| cfa-Let-7-P2b2-5p | Transcriptional regulation (GO:0006357, 0006366)<br>Appendage and limb development (GO:0030326, 0035113, 0035107, ...) |
|  | Transcriptional regulation (GO:0003700, 0000981, 0140110, ...)<br>Post-transcriptional regulation (GO:0030371, 0000900)<br>CCR1 chemokine receptor binding (GO:0031726) |
|  | Cancer (hsa05206, hsa05205, hsa05214) |
| cfa-Mir-1-P3-3p | Chromatin modification (GO:0018027, 0051567, 0016570, ...) |
|  | Chromatin modification (GO:0046974, 0046976, 0018024, ...)<br>Transcriptional regulation (GO:1990841, 0070742, 0140110, ...)<br>Cytoskeletal protein binding (GO:0008092) |
| cfa-Mir-423-3p and cfa-Mir-155-5p see above |  |

**Supplementary Table 1d.** Summarized gene ontology enrichments of targets of historic skin specific microRNA

| microRNA | Enriched terms |
| --- | --- |
| cfa-Mir-203-v1-3p<br>(cfa-Mir-203-v2-3p is similar) | Response to stimulus (GO:0071495, 0009719)<br>Epithelium and blood vessel development (GO:0060429, 0001568)<br>Positive regulation of smooth muscle cell migration (GO:0014911)<br>Cell death (GO:0008219, 0012501, 0006915) |
|  | Transcriptional regulation (GO:0070888, 0000981, 0003700, ...)<br>Growth factor receptors (GO:0008083, 0030618) |
|  | Synapse (hsa04727)<br>Growth factor (hsa04350, hsa04935, hsa01521) |
| cfa-Mir-146-P2-5p | Ectodermal development (GO:0007439, 0048611, 0010668, ...)<br>Transcriptional regulation (GO:0051254, 0045893, 1903508, ...)<br>Appendage and limb development (GO:0030326, 0035113) |
|  | Transcriptional regulation (GO:0001228, 0140110, 0003700, ...)<br>Synapses (GO:0000149, 0019905) |
| cfa-Mir-27-P1-3p | Protein modification (GO:0006464, 0036211) |
|  | Transcriptional regulation (GO:0001228, 0140110, 0003700)<br>Ion binding (GO:0043167, 0046872, 0043169) |
| cfa-Mir-205-P1-5p and cfa-Mir-378-3p see above |  |

